# Acute and chronic toxicity of imidacloprid in the pollinator fly, *Eristalis tenax*

**DOI:** 10.1101/2022.02.18.481021

**Authors:** Nicolas Nagloo, Elisa Rigosi, David C. O’Carroll

**Affiliations:** Department of Biology Lund University, Lund, Sweden

**Keywords:** Imidacloprid, toxicological assay, LD50, chronic exposure, acute exposure, *Eristalis*

## Abstract

Imidacloprid is a neonicotinoid neurotoxin that remains the most used insecticide worldwide. It persists in the environment long after the initial application resulting in chronic exposure to non-target insects. To accurately map the dose-dependent effects of these exposures across taxa, toxicological assays need to assess various modes of exposure across relevant indicator species. However, due to the difficulty of these experiments, contact bioassays are frequently used to quantify dose, and dipterans remain underrepresented. Here, we developed a novel naturalistic feeding bioassay to precisely measure imidacloprid ingestion and its toxicity for acute and chronic exposures in a dipteran pollinator, *Eristalis tenax*. Flies which ingested imidacloprid dosages lower than 12.1 ng/mg all showed consistent intake volumes and learned improved feeding efficiency over successive feeding sessions. In contrast, at doses of 12.1 ng/mg and higher flies had a rapid onset of severe locomotive impairment which prevented them from completing the feeding task. Neither probability of survival nor severe locomotive impairment were significantly higher than the control group until doses of 1.43 ng/mg or higher were reached. We were unable to measure a median lethal dose for acute exposure (72 hours) due to flies possessing a relatively high tolerance for imidacloprid. However, with chronic exposure (18 days), mortality went up and an LD50 of 0.41 ng/mg was estimated. Severe locomotive impairment tended to occur earlier and at lower dosages than lethality, with ED50s of 0.17 ng/mg and 7.82 ng/mg for acute and chronic exposure, respectively. Although the adult *Eristalis* is a honeybee mimic, it possesses a much higher tolerance to this toxin than its model. The similarity in the LD50 to other dipterans such as the fruitfly and the housefly suggests that there may be a phylogenetic component to pesticide tolerance that needs to be further investigated. The absence of obvious adverse effects at sublethal dosages also underscores a need to develop better tools for quantifying animal behaviour to evaluate the impact of insecticides on foraging efficiency in economically important species.

## 1 Introduction

It is widely accepted that pesticides are one of the leading factors driving biodiversity loss, especially in insect populations (Geiger et al., 2010; Hallmann et al., 2017; Sánchez-Bayo and Wyckhuys, 2019). Despite concerns over such declines, regulatory authorities have been slow to develop effective mechanisms to assess risk on diverse insect groups. In Europe, for example, longstanding environmental risk assessment (ERA) guidelines of the European Food Safety Authority (EFSA), define risk posed by a pesticide as the product of pesticide toxicity and the likelihood of exposure in field settings (Condalfi et al., 2000). Yet at the first level of ERAs (Tier 1) risks to insects and other arthropods are extrapolated from data gathered on just two indicator species, a parasitoid wasp *Aphidius rhopalosiphi*, and a predatory mite, *Typhlodromus pyri* (Condalfi et al., 2000). Both species are beneficial to crops, and are easily reared in laboratory settings, allowing for larger sample size, and facilitating the measure of impacts on reproduction. They are also believed to be suitably sensitive to pesticides (Mead-Briggs et al., 2000; Grimm et al., 2001). However, more recently concerns over impacts on the pollinating role of bees has resulted in the creation of a separate guidance document (EFSA, 2013) which requires additional tests in honeybees, *Apis mellifera*, and bumblebees, *Bombus terrestris*, to better understand the impact of pesticides on bees. Despite this welcome expansion, the number of arthropod species range between 5 and 10 million (Ødegaard, 2008) and they occupy a wide range of ecological niches, making them susceptible to a variety of exposure routes not explored by standardized tests. In addition to insufficient biodiversity coverage, recent reviews of these guidance documents by panels of the EFSA have highlighted the need for the inclusions of tests which examine the impact of repeated exposure over longer periods of time (EFSA, 2015; 2018). This has produced an increasing body of research which demonstrates that even if organisms survive initial exposure, sublethal dosages can have important effects on learning performance, neurophysiology and behaviors ranging from feeding to navigation (Nauen, 1995; Decourtye et al., 2003; Easton and Goulson, 2013; Laycock and Cresswell, 2013; Tappert et al., 2017; Parkinson and Gray, 2019).

To determine suitable toxicity endpoints for lethal and sublethal exposures, precise measures of pesticide intake and its subsequent effects are necessary. However, most toxicological studies measure the concentration of pesticide that is lethal to 50% (LC50) of the population without regard for individual intake (dose) (Mizell and Sconyers, 1992; Overmyer et al., 2005; Stoughton et al., 2008; Cavallaro et al., 2017). In studies which measure individual dosage (LD50), insecticides are typically administered through artificially applied contact due to the ease of dose measurement (Wang et al., 2002; Da Costa et al., 2015; Soares et al., 2015; Tappert et al., 2017). However, this overlooks alternative and more natural routes of exposure, such as oral intake where the pesticide may be more toxic (Tharp et al., 2000; Nauen et al., 2001; Marletto et al., 2003).

Insecticide use across the globe has intensified since the 1940s, with cholinergic pesticides such as neonicotinoids used in 120 countries and across 450 crops (Simon-Delso et al., 2015; Bakker et al., 2020). Imidacloprid was the first introduced neonicotinoid and in 2010, estimated production ranged up to 20,000 tons (Kagabu, 2011). Despite the banning of imidacloprid, thiamethoxam, and clothianidin in May 2018 due to their adverse effects on bees (EFSA, 2018), many Member States still provide emergency authorization for the use of these banned substances. These neonicotinoids affect a range of non-target species and imidacloprid tends to be more toxic through oral intake than upon contact in bees, bumblebees, and grasshoppers (Suchail et al., 2000; Tharp et al., 2000; Nauen et al., 2001). The orders of magnitude between bee and pest species’ sensitivity to imidacloprid suggest that extrapolation of toxicity endpoints should be done carefully, as imidacloprid sensitivity can vary greatly across taxa (Lagadic et al., 1993; Suchail et al., 2001; EFSA, 2018).

Flower visiting flies are an ecologically important subset of pollinators. However, under current guidelines, the toxicity of imidacloprid to non-bee arthropods would most likely be derived from toxicity endpoints of *T. pyri* and *A. rhopalosiphi*. Out of the many flower-visiting dipterans, *Eristalis tenax* is one of the most ecologically important non-bee pollinators for both wild flowers and crops (Jarlan et al., 1997; Ssymank et al., 2008; Gervais et al., 2018; Howlett and Gee, 2019; Cook et al., 2020; Doyle et al., 2020; Baumann et al., 2021). Previous studies reveal some tolerance to the neonicotinoid thiamethoxam in the aquatic larvae of *Eristalis* with concentrations greater than 500 µg/L required to produce significant mortality (Basley et al., 2018). Significant neurophysiological impacts of imidacloprid which alter motion detection have also been observed in *Eristalis* (Rigosi and O’Carroll, 2021). However, the dose-dependent lethal and sublethal impacts of imidacloprid remain unclear.

Here we examined the lethal and sublethal impacts of imidacloprid ingestion in this ecologically important fly. Our controlled feeding experiments allowed us to overcome some limitations of previous LC50 and LD50 studies in other species by targeting a relevant natural oral exposure route and precisely measuring individual intake. Our results suggest that *Eristalis* is at least an order of magnitude more tolerant to imidacloprid than bees. Low to intermediate doses have no negative impacts on survival and feeding volume over time. In contrast, at very high doses of 1.43 ng/mg or higher, imidacloprid impairs mobility and becomes lethal, especially with chronic exposure.

## 2 Materials and methods

### 2.1 Insect husbandry

Drone flies, *Eristalis tenax*, pupae were obtained from POLYFLY SL (Spain). After hatching they were allowed to mature for 5-6 days before healthy males were selected, weighed, and isolated for 24hrs prior to the start of feeding trials. On average, we used 50 males per group with a minimum of 29 and a maximum of 58 individuals per treatment group (Table 1). Flies were kept in good condition by closely matching protocols outlined by Nicholas et al (2018) for long-term maintenance of this species. Briefly, hoverflies were kept at 11 °C in a fridge and participated in feeding sessions every three to four days. On feeding days, flies were kept at ∼25 °C for four to six hours before being returned to the fridge to await the next feeding session. This regimen has been shown to keep animals healthy in captivity for periods of up to one year (Nicholas et al., 2018). *Eristalis spp*. persist in the wild even during the late autumn and winter in northern Europe (Stubbs and Falk, 1983). Feeding activity in larger hoverflies such as *Eristalis* occur at all temperatures above 11 °C (Gilbert, 1985) suggesting that the fridge temperature (11 °C) used in this study reflects conditions experienced in the wild.

**Table 1:**
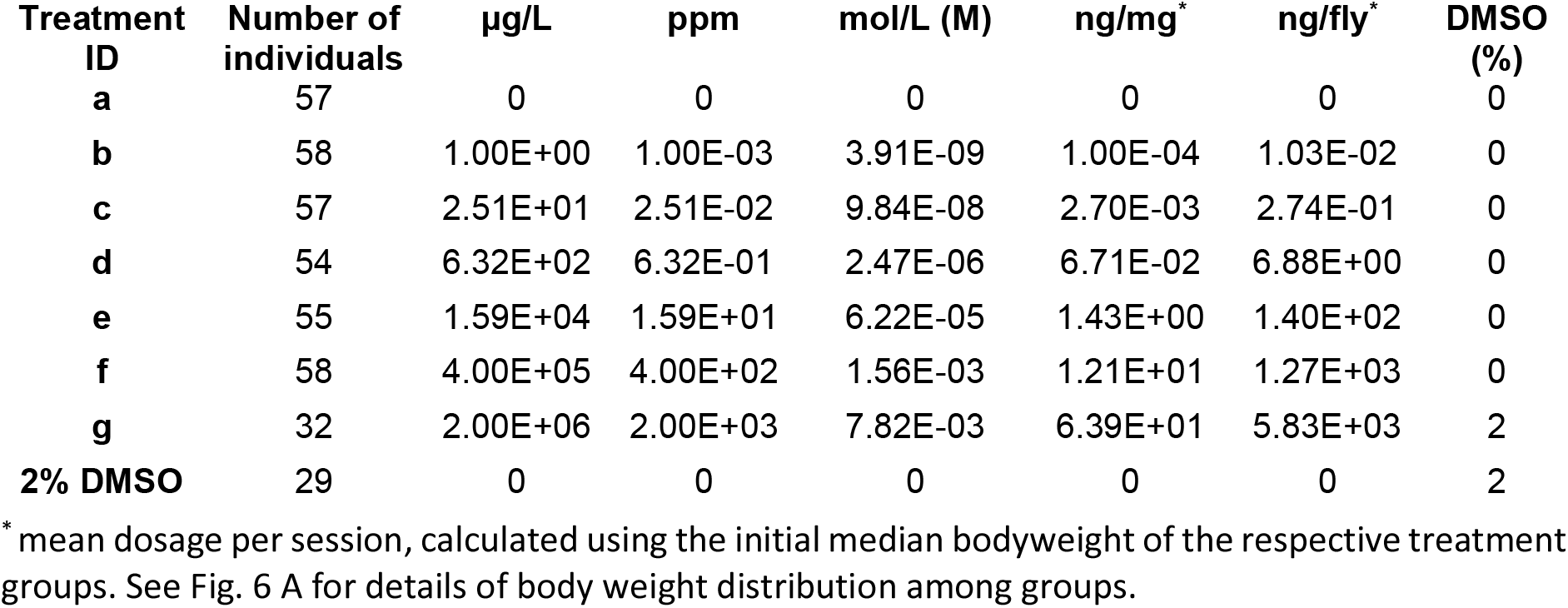
Imidacloprid concentrations and number of individuals per group

### 2.2 Chemicals and dilutions

Due to the poor solubility of imidacloprid in water, most toxicology studies to date have dissolved imidacloprid either in dimethyl sulfoxide (DMSO) or acetone. However, due to the adverse effects of DMSO in *Eristalis* and *Drosophila melanogaster* (Cvetković et al., 2015; Rigosi and O’Carroll, 2021), we attempted to avoid the use of a non-aqueous solvent where possible. We dissolved analytical grade imidacloprid (CAS Number: 138261-41-3, Sigma Aldrich, Sweden) in a solution of 10% raw sugar in 0.01M Phosphate Buffer to achieve logarithmically spaced concentrations of imidacloprid ranging from µg/L to 2 g/L (Table 1) until we exceeded the solubility limit of imidacloprid in water (610 mg/L) (Tomlin, 2006). Only our highest concentration (Treatment g, Table 1) was dissolved in a combination of DMSO and sucrose solution to reach 2 g/L of imidacloprid in solution. We added a 2% DMSO vehicle treatment to control for DMSO effects in the experiment.

### 2.3 Feeding experiments

The feeding trials lasted for 15 days with feeding events occurring every three to four days for a total of five exposures to control or imidacloprid solutions over 15 days. Feeding occurred in chambers dedicated to a specific treatment to avoid cross-contamination. These consisted of an inverted cylindrical plastic container (Outer diameter (OD): 52 mm, 125 ml, VWR International, USA) and a custom-made 3D printed white flower with a yellow centre mounted on the wall of the container (Fig. 1). A hole in the centre of the flower allows a non-heparinised 75 µl micro-capillary (Hirschmann, Germany) to be introduced into the feeding chamber at an angle of 15° to give the fly access to the feeding solution of choice (Fig. 1). At the beginning of each feeding trial, a mark was made at the initial meniscus of the feeding solution on the glass capillary and at a target volume 10 µl below that mark. The glass capillary was removed once an individual had either consumed 10 µl or reached a “time out” set at 90 minutes for the first feeding session and 60 minutes for subsequent sessions. The time taken to feed was then recorded as when either 10 µl had been consumed or time out was reached. In a small number of trials, the fly consumed food so rapidly that the level passed the 10 µl mark before the experimenter was able to terminate access. The actual volume consumed was, however, then recorded as the difference between the initial mark and the final meniscus. This protocol minimises insect handling, allows the monitoring of feeding behaviour and allows for control over the dose administered to the animal. After the feeding session, flies were placed in a separate chamber (OD: 33mm, 60 ml, VWR International, USA) where they were provided with 20 ul of honeyed water (Acacia honey, COOP, Sweden, 1 drop in 1ml of distilled water) and a grain of organic pollen (Rawpowder, Sweden) for a total time of four hours at RT (including time in the feeding chamber). Flies were then returned to the fridge at 11 °C in individual containers (OD: 33mm, 60 ml, VWR international, USA) with 50 µl of distilled water on piece of filter paper.

**Fig. 1:**
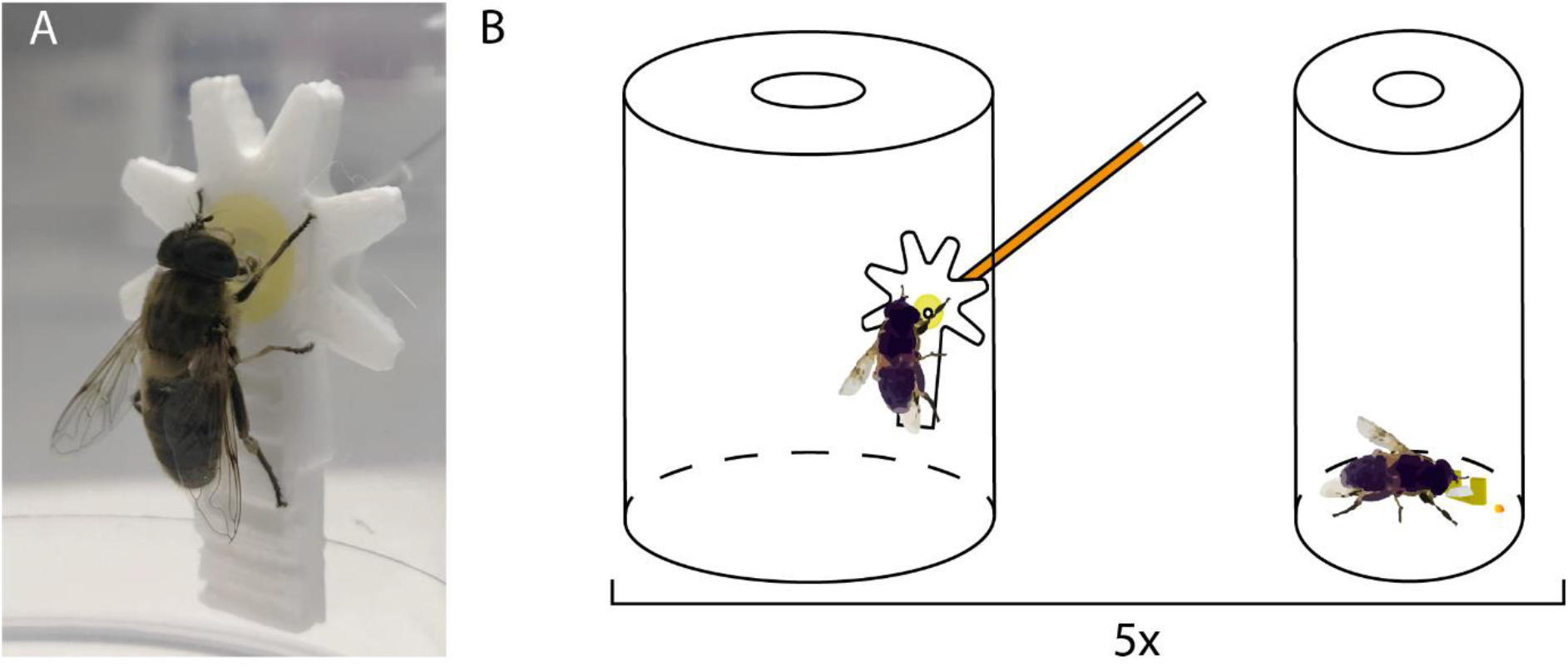
A new feeding paradigm to control pesticide exposure at individual level in the pollinator fly, *Eristalis tenax*. A) An example of a fly feeding from a capillary filled with imidacloprid and sucrose solution and inserted in the center of a 3D printed flower. B) Each fly was exposed 5 times to a solution of imidacloprid (or vehicle) that was accessible from a 3D printed flower (A) in their feeding chamber (125 mL container). The meniscus of the solution (false colored here) was measured before and after ingestion to quantify the ingested volume per individual. After exposure, flies were left in 60 ml container up to 4 hours at RT with *ad libitum* access to honey water and a grain of pollen.

### 2.4 Insect status observations

Fly status was observed and recorded every one to two days for 18 days from the start of the feeding trials. Individual status at each point was classified into three broad categories: ‘mobile’, ‘severe locomotive impairment’ or ‘dead’. Severe locomotive impairment was recorded when a fly ended up on its back and was unable to get up. Flies able to move around the chamber and climb were classified as mobile, even if they exhibited some obvious signs of motor impairment. Death was recorded when there was a complete lack of response to tactile stimuli more than 30 minutes after the last feeding session.

### 2.5 Acute and chronic dose response curves

Cumulative status observations collected at 72 hrs (3 days after the first feeding session) and 432 hrs (18 days total, i.e. 3 days after the 5^th^ feeding session) were used to assess acute and chronic impacts, respectively. Acute observations thus assessed the impact of a single exposure while chronic observations assessed the impact of 5 successive exposures over 15 days. The dosage of imidacloprid administered to an individual was calculated as the mean amount ingested across all feeding sessions which was then divided by the median body weight of the treatment group, measured 24 hrs prior to the initial feeding session to provide a mass-specific intake (ng of imidacloprid/mg of bodyweight). Observations from all feeding trials were pooled to determine the lethality/ locomotive impairment induced by imidacloprid ingestion across dosages at 72 hrs and 18 days. A five parameter logistic regression curve (Cardillo, 2021) which optimises minimum asymptote, Hill’s slope, inflection point, maximum asymptote and asymmetry was fitted to the data using a non-linear least-squares method (Levenberg-Marquardt). This allowed the estimation of median lethal dose (LD50) and median effect dose (ED50) when examining lethality and impairment/death respectively.

### 2.6 Effect size on feeding behaviour

We also assessed sublethal impacts of imidacloprid on feeding behaviour. The effect of a feeding session on the volume consumed and the time taken to consume the standard 10 µl was quantified by taking the difference of means between successive sessions, standardised by the weighted pooled standard deviation (Equation 1) to obtain Hedges’ g as a measure of the effect size (Equation 2). We first computed a weighted pooled standard deviation *S*_*p*_ as

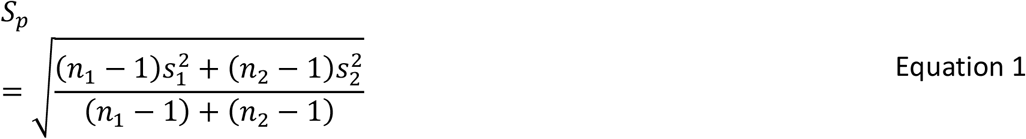

where *n*_1_ and *n*_2_ are the sample sizes of their respective groups, and *s*_1_ and *s*_2_ are the standard deviation of their respective groups. We calculated effect size *g* as

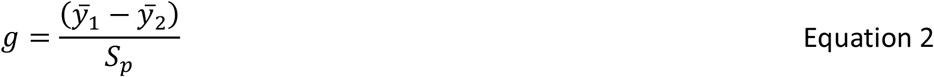

where 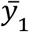 and 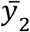 are the means of their respective groups, with *S*_*p*_ being the previously calculated pooled standard deviation (Hedges et al., 1985). Effect size was then bias corrected (Equation 3) for combined sample sizes less than 40 as

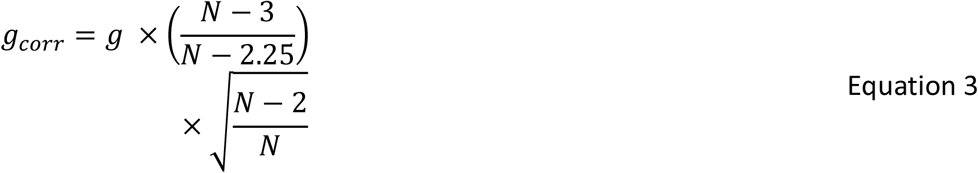

where *g*_*corr*_ is the bias corrected effect size, *g* is the effect size and N is the combined group sample size (Durlak, 2009). 95% confidence intervals were then obtained using the percentile bootstrap method with 1000 iterations (Efron, 1982). Calculations and bootstrapping were conducted in custom-written functions in Matlab (Mathworks, MA, USA).

### 2.7 Impact of imidacloprid on likelihood of survival and impairment

Probability of survival or impairment and death over time were calculated using the Kaplan-Meier method from the *Surv* module of the survival package in R (R Core Team, 2021). Due to the possibility of non-proportional hazard across treatments, an alternative to the popular log-rank test, the Peto&Peto modification of the Gehan Wilcoxon test is used to compare survival curves (Gehan, 1965; Peto and Peto, 1972). The *survdiff* module of the survival package in R was used for statistical comparisons (R Core Team, 2021). Due to the increased likelihood of false positives inherent to multiple pairwise comparisons between control and treatments, the p-value was adjusted using the Bonferroni method using the *p*.*adjust* module from the stat package in R (R Core Team, 2021).

## 3 Results

Our novel feeding system promoted active and naturalistic feeding behaviour by flies as well as providing precise measurements of imidacloprid intake. Flies readily learned to feed from the chambers during their first feeding trial. The yellow centre of the 3D printed flower (Figure 1) acts as a guiding cue and is known to elicit an unconditioned proboscis extension reflex in *Eristalis spp*., more so than in other hoverflies (Lunau et al., 2018). In initial trials with sugar solution, we found that an angle for the feeder tube of 15° is sufficient to balance capillary force with gravitational pull. Individual flies given *ad libitum* access to this feeder would readily consume 3-6 times our target volume of 10 µl of feeding solution, confirming that any reduction in intake was more likely to be due to toxic effects of the treatment than an ability to learn the feeding task. In addition to providing the means to control dosage, our feeder thus allows insights into the dose-dependent impacts of imidacloprid on feeding behaviour that would be unachievable by traditional glass plate toxicological assays (Blümel et al., 2000; Mead-Briggs et al., 2000).

### 3.1 Effects of successive feeding sessions on feeding amount and time

Figure 2 shows the treatment volume consumed by individuals and the proportion of feeding individuals in five successive feeding sessions. For concentrations up to 6.32 × 10^2^ µg/L, the consumed volume per individual was close to the 10 µl target across all feeding sessions (**Fig. 2 A**). The number and percentage of live flies feeding also remained relatively constant across feeding sessions, suggesting minimal impacts of the pesticide on feeding behaviour at concentrations of up to 6.32 × 10^2^ µg/L (**Fig. 2 B**). While the median consumed volume remains close to 10 µl at an imidacloprid concentration of 1.59 × 10^4^ µg/L, greater variation is observed around the median suggesting that individual flies were differentially affected at this concentration (**Fig. 2 A**). Furthermore, the proportion of flies feeding across feeding sessions declined suggesting that their ability to feed began to deteriorate due to repeated imidacloprid ingestion (**Fig. 2 B**). At higher concentrations of 4.00 × 10^5^ µg/L and above, the median consumed volume dropped to between 0 and 3 µl (**Fig. 2 A**). Ingestion of imidacloprid at such high concentrations, even at such small volumes was accompanied by a violent burst of wing beats and complete paralysis within 30 seconds of ingestion. There was also a sharp drop in the number and proportion of live individuals feeding by the second feeding session suggesting that they had not yet recovered from the acute impacts observed during the initial feeding session (**Fig. 2 B**). This sharp decline in the second feeding session was followed by a gradual increase in the proportion of flies feeding in subsequent sessions (Fig. 2B). This implies that surviving flies partially recovered their feeding ability after the initial severe acute impact of exposure to high concentrations of imidacloprid.

**Fig. 2:**
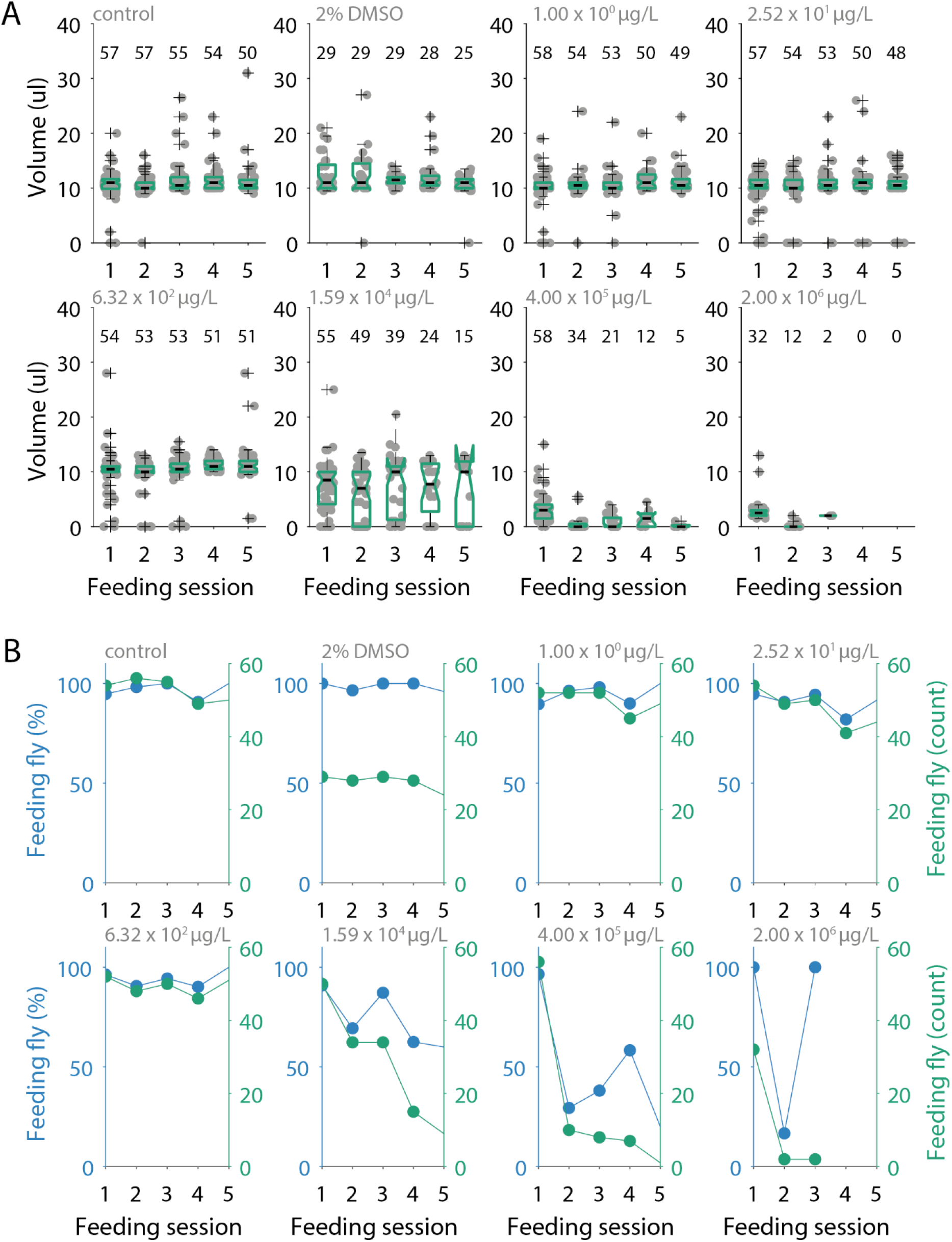
Dose-dependent effects of imidacloprid on consumption volume and number of feeding individuals over time. A) Volume consumed within feeding sessions over time for each treatment. B) Number and proportion of feeding flies over time for each treatment. The volume consumed by individual flies as well as the number and proportion of flies remained constant throughout the experiment for flies exposed to imidacloprid concentrations of up to 6.32 × 10^2^ µg/L in their diet. Above that, the number and proportions of flies feeding started decreasing over time with smaller and more variable volumes consumed by individual flies. In panel A, grey filled circles indicate individual data points plotted with horizontal jitter to increase data clarity. This is overlayed with boxplots with box limits denoting the 25^th^ and 75^th^ percentile. Area between the whiskers span ± 2.7 standard deviations, anything outside is drawn as an outlier denoted by a black cross. Notches on the boxplot indicate 95% confidence interval of the median. The number above each boxplot denotes the number of flies included in that group.

Figure 3 shows the effect of repeated exposure to imidacloprid on feeding volumes. The drop in the number of feeding flies in treatment groups exposed to high imidacloprid concentrations (> 4.00 × 10^5^ µg/L) after the initial exposure prevents us from reliably computing the effect size for affected groups. However, across all remaining groups, the average absolute effect size of repeated exposure in feeding volume was small (0.18) and mostly not significant. Two exceptions were observed with flies consuming significantly less in the second feeding session of the control group and the fifth feeding session of the DMSO control group (**Fig. 3**). Overall, this suggests that there is no cumulative effect of imidacloprid exposure on the volume that flies consume across all concentrations.

**Fig. 3:**
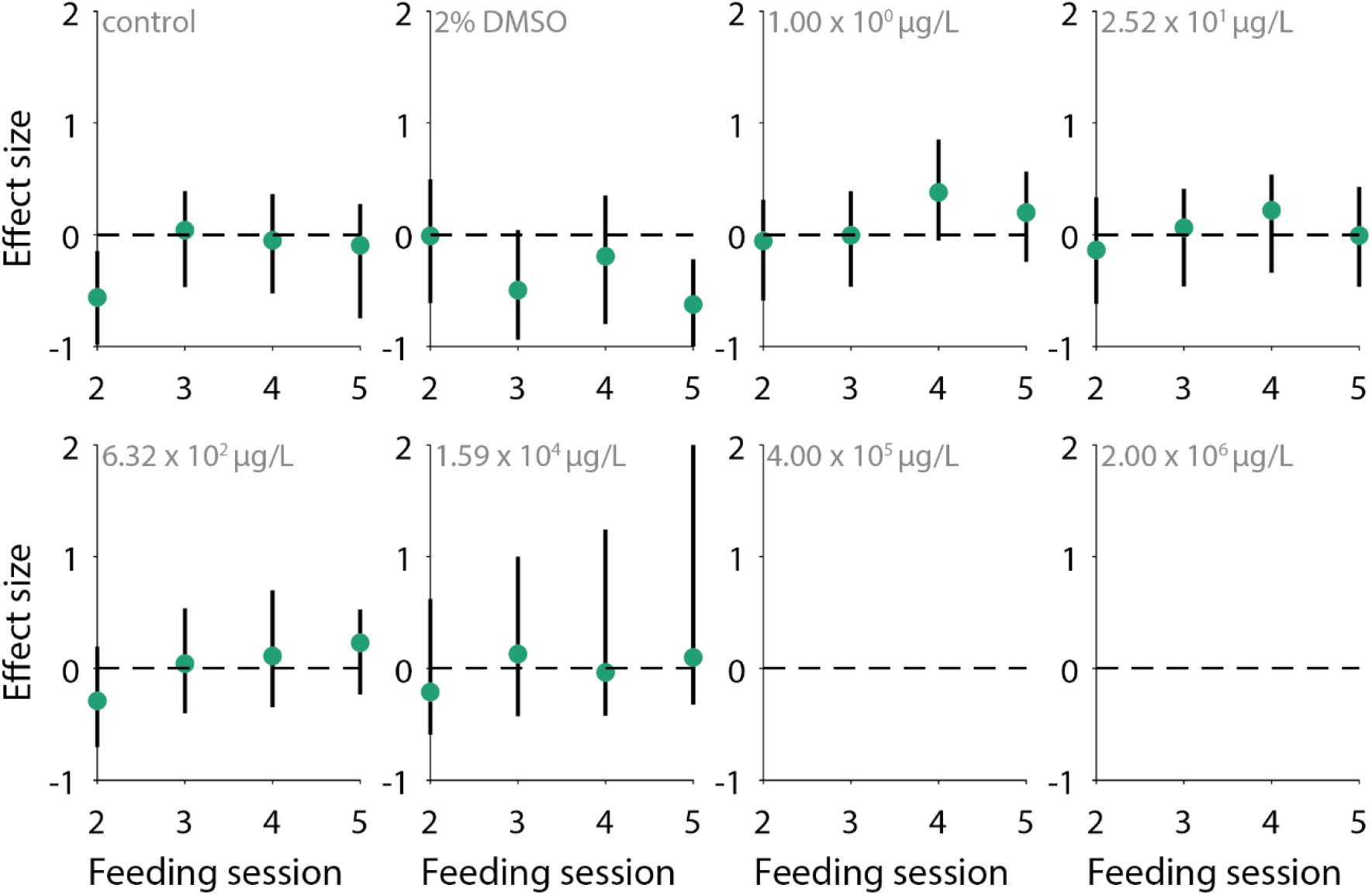
The effect of successive imidacloprid exposures on consumption volume relative to the first feeding session. Effect size remained small with an average of 0.18 ± 0.18 across all feeding sessions and imidacloprid treatments. Volume consumed across feeding sessions were typically not significantly different from the initial feeding session for any given concentration of imidacloprid. Green circles indicate the Hedges’ g effect size of a feeding session relative to the initial feeding session for any given treatment. Black vertical bars indicate the 95% confidence interval of effect sizes while horizontal black dashed lines indicate zero where no effect is observed. If the confidence interval crosses the horizontal dashed line, effects are considered not significant. Effect sizes are interpreted as small (0.2), medium (0.5) or large (0.8).

Figure 4A shows the time taken for individual flies to consume 10 µl, while figure 4B shows percentage of feeding flies which took longer than 60 minutes to feed (‘time out’). The time taken for individual flies to consume 10 µl decreased over successive feeding sessions (**Fig. 4 A**) with lower percentages of feeding flies taking more than 60 minutes to feed in subsequent feeding sessions (**Fig. 4 B**). Figure 5 shows the effect of successive feeding sessions on feeding time (relative to the first). Flies typically took significantly less time to feed (i.e., the upper 95% confidence interval of the effect size was below zero) than in their initial naive feeding session by the 3^rd^/4^th^ feeding session (**Fig. 5**). This suggests that the flies learned how to feed more efficiently inside their feeding chambers. This was seen for all concentrations of 6.32 × 10^2^ µg/L and lower (**Fig. 4A, Fig. 5**). The effect size after five feeding sessions was largest in the control group (Hedges’ g ± 95%CI = -1.13, [-0.71, -1.58]) and smallest at the highest imidacloprid concentration (Hedges’ g = -0.25, [0.32, -0.80]) suggesting that this learning effect was weaker following exposure to high concentrations of imidacloprid (**Fig. 5**). As described earlier, at the two highest concentrations, most flies suffered from severe mobility impairment soon after ingestion and were unable to consume 10 µl within the allotted time with between 94% and 100% of the flies timing out (**Fig. 4 B**). Nevertheless, it is worth noting that as long as flies were not exposed to high enough concentrations of imidacloprid (> 4.00 × 10^5^ µg/L) to paralyse them upon initial ingestion, median consumption volumes remained close to the 10 µl across feeding sessions and feeding times still improved over time (**Fig. 2 A, Fig. 4 A**). This strongly suggests that imidacloprid does not have a direct antifeedant effect at intermediate doses as they will still feed if they are able to reach their food source.

**Fig. 4:**
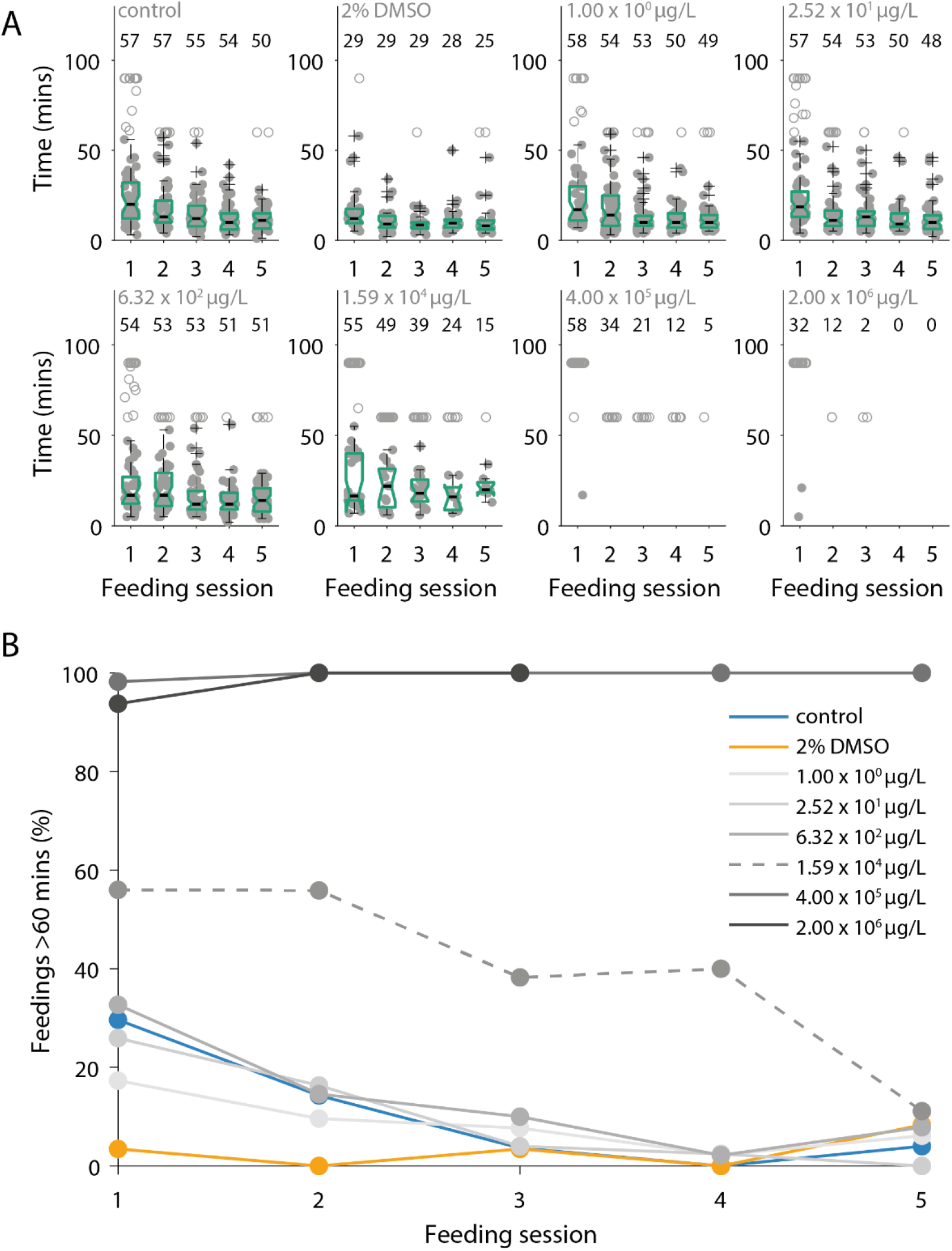
Dose-dependent effect of imidacloprid on feeding time in *Eristalis*. A) Time taken to find and consume 10 µl of solution across feeding sessions for each treatment. B) The percentage of feeding individuals which take longer than 60 minutes to feed for each treatment. Flies become faster at feeding over time which results in less flies running out of time to complete their feeding task. In panel A, grey filled circles indicate individual data points plotted with horizontal jitter to increase data clarity. This is overlayed with boxplots with box limits denoting the 25^th^ and 75^th^ percentile. Area between the whiskers span ± 2.7 standard deviations, anything outside is drawn as an outlier denoted by a black cross. Notches on the boxplot indicate 95% confidence interval of the median. The number above each boxplot denotes the number of flies included in that group. In panel B, the in-figure legend shows the concentration of imidacloprid in each treatment group.

**Fig. 5:**
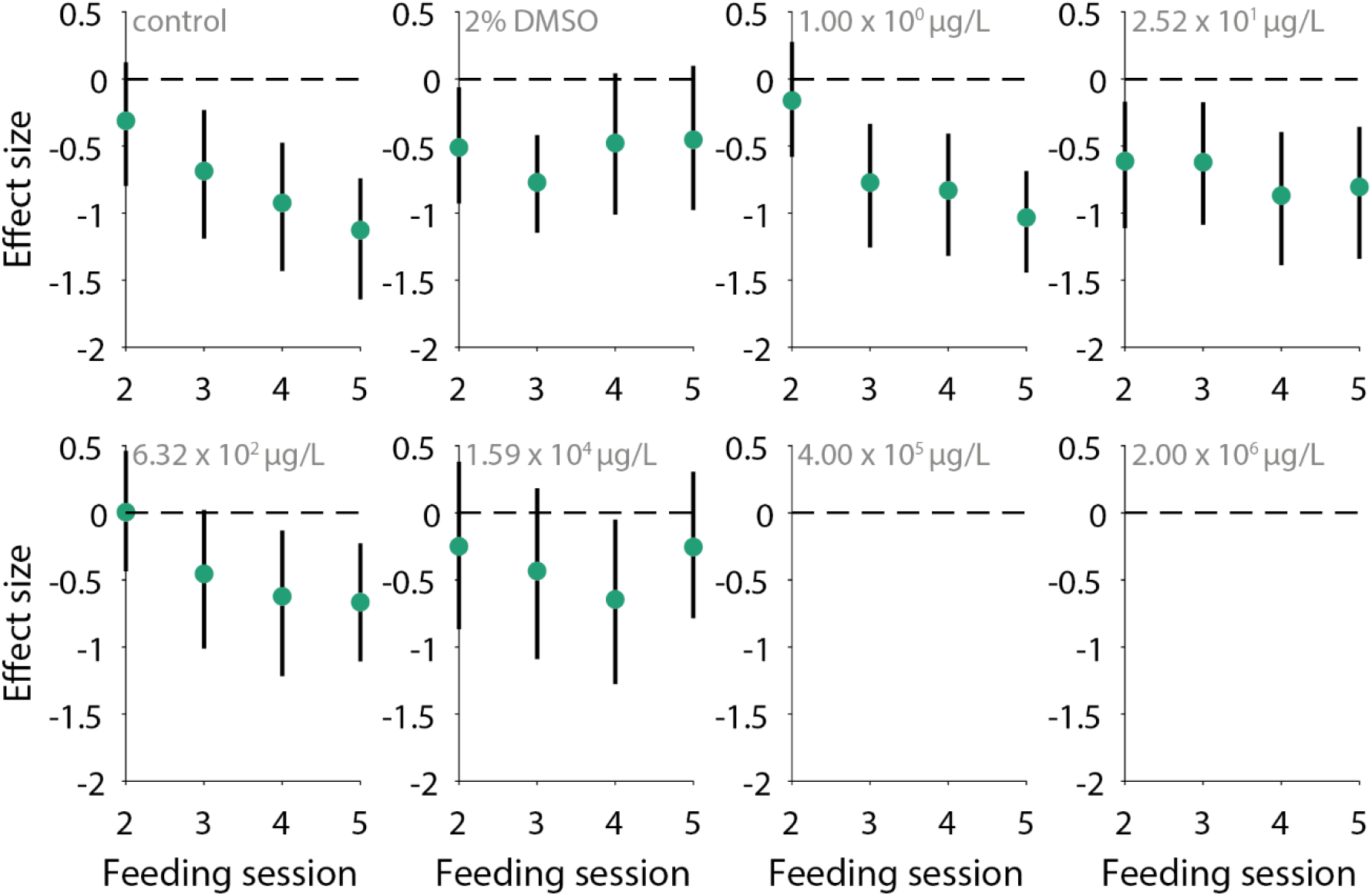
The effect of successive imidacloprid exposures on feeding time relative to the first feeding session. Successive feeding sessions had a medium to large effect on time taken to consume 10ul with an average absolute effect size of 0.60 ± 0.28 (mean ± s.d.) across all dosages and over all feeding sessions. Flies learned how to feed more efficiently and generally became faster than the initial feeding session by the 3^rd^/4^th^ feeding session. Green circles indicate the Hedges’ g effect size of a feeding session relative to the initial feeding session for any given treatment. Black vertical bars indicate the 95% confidence interval of effect sizes while horizontal black dashed lines indicate zero where no effect is observed. If the confidence interval crosses the horizontal dashed line, effects are considered not significant. Effect sizes are interpreted as small (0.2), medium (0.5) or large (0.8).

Median pre-experimental body mass across treatment groups ranged from 90 mg to 105 mg with an average of 102 ± 16 mg (**Fig. 6 A**). Imidacloprid intake of individual flies in a single feeding session thus ranged from 2.0 × 10^−3^ ng to 2.6 × 10^4^ ng across feeding sessions and treatments (**Fig. 6 B**). Despite paralysis induced upon ingestion at the two highest concentrations, the relatively straight line in the log-linear plot in Figure 6 C shows that the median imidacloprid dosage kept increasing exponentially across treatment groups (**Fig. 6 C**).

**Fig. 6:**
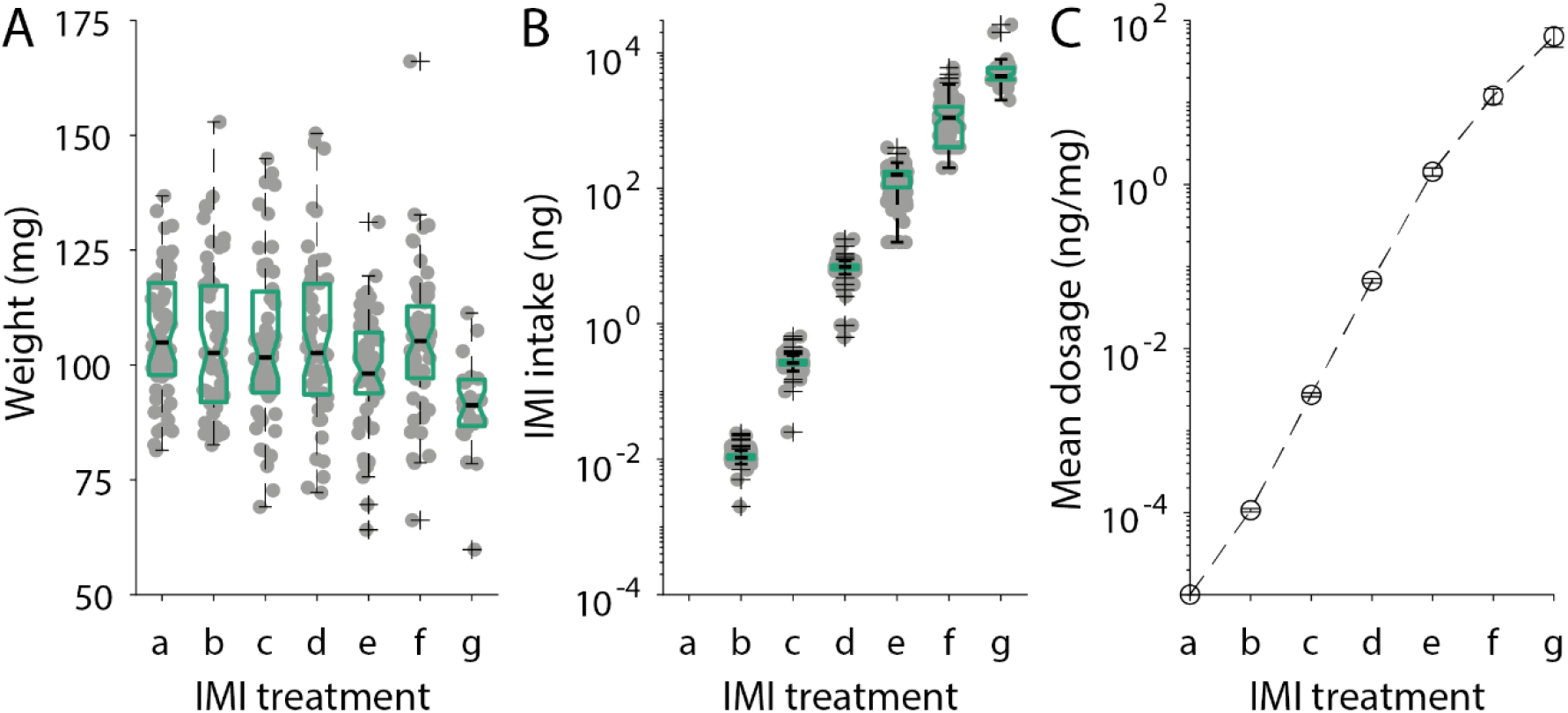
Intake of imidacloprid across treatments and correction for body weight. A) Insect body weight before imidacloprid exposure for each treatment. B) Distribution of net imidacloprid intake across treatments. C) Mean dosage achieved across feeding sessions across treatments standardised to the median initial bodyweight of respective treatments. Despite severe mobility impairment, imidacloprid intake does not decrease and the effective dosage in ng/mg of bodyweight continues to increase exponentially across treatments. For boxplots in panel A and B, box limits denote the 25^th^ and 75^th^ percentile. Area between the whiskers span ± 2.7 standard deviations, anything outside is drawn as an outlier denoted by a black cross. Notches on the boxplot indicate 95% confidence interval of the median. Error bars in panel C indicate the 95% confidence interval of the mean. Labels a-g stand for the concentrations of imidacloprid, refer to Table 1.

### 3.2 Lethality and severe impairment induced by imidacloprid over time

Figure 7 shows the dose-dependent probability of survival and impairment/death with time and the estimation of LD50 and ED50 for acute and chronic exposure to imidacloprid. The probability of survival after 18 days when ingesting the control solution was 0.84 [0.75 0.94] (**Fig. 7 A**). There were no differences in survival rates between control and all treatments equal to or lower than 6.32 × 10^2^ µg/L as well as the 2% DMSO vehicle (**Table 2**). It was not possible to calculate a median survival time for these treatments as survival probabilities remained above 0.5 throughout the experiment, confirming the efficacy of our maintenance protocol (**Fig. 7A**). In contrast, survival at concentrations of imidacloprid above 6.32 × 10^2^ µg/L (treatment d in Tables and Fig. 6), was significantly different from control (**Table 2**) with increasingly short median survival times (days) [lower 95%CI, upper 95%CI] of 9 [8,13], 5 [2,7], and 2 [1,5] for the 3 highest dose treatments (e, f, and g, respectively). Survival probability shortly after the first exposure remained high such that it was impossible to calculate a median lethal dosage (LD50) for a single acute exposure after 72hrs (**Fig. 7 B**). However, with chronic exposure, the decreased survival probability enabled the calculation of the median LD50 as 0.41 ng/mg.

**Table 2:**
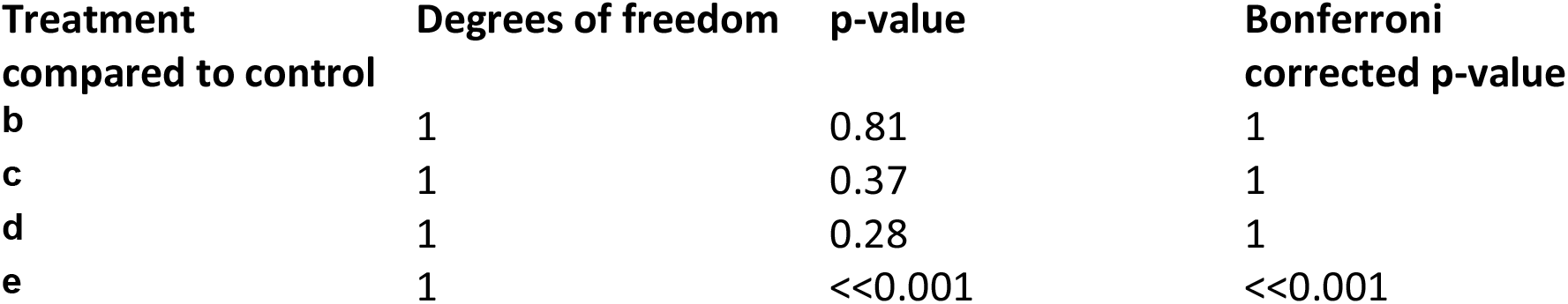
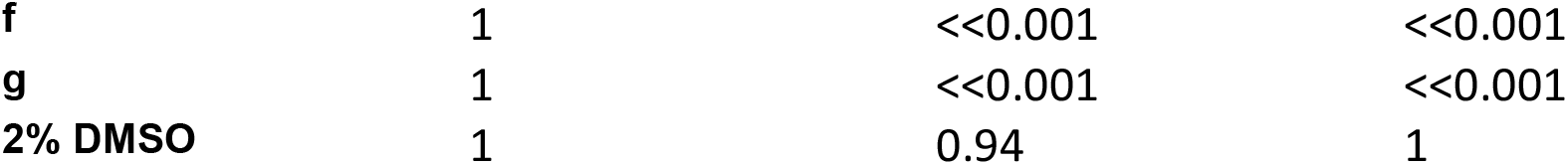
Comparisons of survival curves to control using a weighted log rank test

**Fig. 7:**
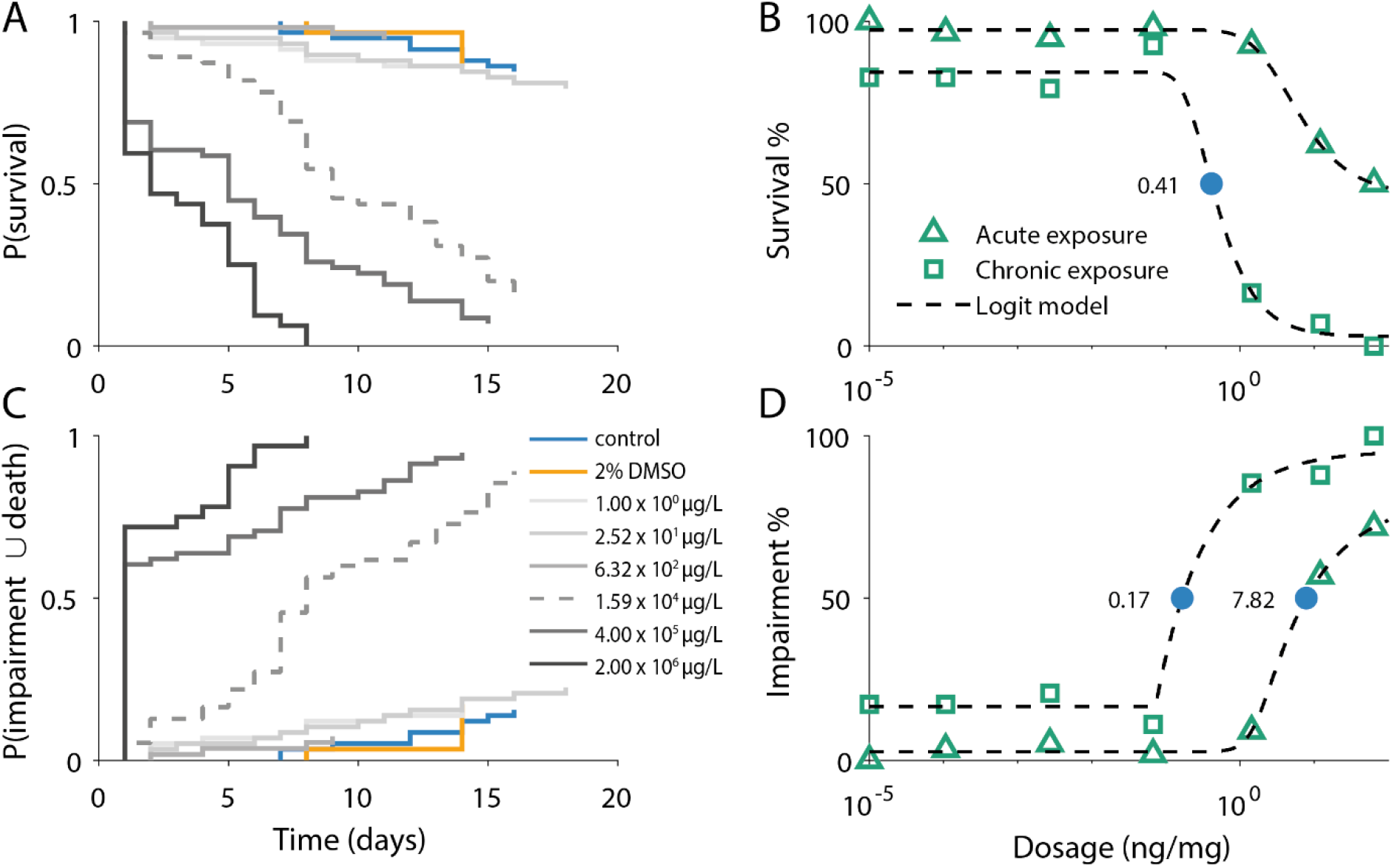
Toxicity of imidacloprid over time and across dosages. A) Survival probability over time for each treatment. B) Survival percentages across dosages for acute and chronic exposure. C) The probability of impairment or death throughout the experiment across dosages. D) The percentages of individuals either impaired or dead throughout the experiment across dosages. Probability of survival remains high with low probabilities of impairment or death throughout the experiment for vehicle controls and at dosages of 6.71 × 10^−2^ ng/mg and lower. Legend values are presented in ng of imidacloprid per mg of bodyweight (ng/mg) in panel A and C. Green triangles represent responses to a single exposure after 72 hrs (acute), green squares represent responses to five exposures after 18 days (chronic) in panel B and D. The blue circle represents the LD 50 in panel B and the ED50 in panel D.

Similarly, the combined probability of impairment and death did not differ significantly from control at concentrations equal to or lower than 6.32 × 10^2^ µg/L or for the 2% DMSO vehicle (**Table 3**) but were significantly higher for all concentrations above 6.32 × 10^2^ µg/L (**Table 3**). For the three highest doses (treatments e, f and g) median time (days) to impairment/death [±95%CI] were 8 [7,12], 1 [1,5], and 1 [1,1], respectively. The much shorter median times to impairment/death compared to median survival times for the same treatments suggest that although *Eristalis* may not die when first exposed to high concentrations of imidacloprid, fly locomotion is very quickly impaired. Median doses to impairment/death for acute and chronic ingestion of imidacloprid were 7.82 ng/mg and 0.17 ng/mg, respectively.

**Table 3:** Comparisons of cumulative hazard curves for impairment/death to control using a weighted log rank test

| **Treatment**<br>**compared to control** | **Degrees of freedom** | **p-value** | **Bonferroni**<br>**corrected p-value** |
| --- | --- | --- | --- |
| **b** | 1 | 0.88 | 1 |
| **c** | 1 | 0.34 | 1 |
| **d** | 1 | 0.42 | 1 |
| **e** | 1 | <<0.001 | <<0.001 |
| **f** | 1 | <<0.001 | <<0.001 |
| **g** | 1 | <<0.001 | <<0.001 |
| **2% DMSO** | 1 | 0.92 | 1 |

## 4 Discussion

### 4.1 Locomotive impairment and antifeedant effects

Throughout the feeding trials used in this study, naïve flies were able to locate and consume consistent volumes from a glass capillary throughout the five feeding sessions while learning to feed significantly faster during the experiment. This contrasts with some work on bees. Honeybees, for example, will reject sucrose solutions with imidacloprid concentrations of 1.0 × 10^3^ µg/L (Nauen et al., 2001) or higher in a dose-dependent manner. Here we observed *Eristalis* attempt to feed at concentrations of up to 2.0 × 10^6^ µg/L despite this leading to immobilisation. At concentrations of 10 µg/L, cumulative antifeedant effects have also been observed in the bumblebee, *Bombus terrestris* (Thompson et al., 2015). Such antifeedant effects observed in *B. terrestris, Myzus persicae* and *Serangium japonicum* are reversible within 24hrs after exposure to imidacloprid is terminated (Nauen, 1995; He et al., 2012; Thompson et al., 2015). This suggests that, at least at lower doses, imidacloprid is at least not repulsive to the flies (i.e., has no direct antifeedant effect). It is worth noting that exposure to imidacloprid may in some other species, such as in the red fire ant, increase consumption at concentrations of 10 µg/L (Sakamoto and Goka, 2021).

At higher doses it becomes difficult to dissociate potential antifeedant effects from locomotive impairment. Flies exposed to concentrations of 4.0 × 10^5^ µg/L (median dose: 12.1 ng/mg) or higher are immobilised within 30 seconds of ingestion and as a result median consumption falls to no higher than 3 µl (**Fig. 2 A**). Similar knock down effects have been observed in honeybees which have ingested an imidacloprid dose of ∼0.72 ng/mg where they become immobilised, such that sucrose intake drops by 33% (Nauen et al., 2001). Nevertheless, while it remains possible that imidacloprid may become completely repulsive to *Eristalis* at higher concentrations, exposure to such levels is extremely unlikely to occur in field exposure. Imidacloprid concentrations detected in fields where treated seeds are in use ranges between 1-22 µg/kg, 1-10 µg/kg, and 1-11 µg/kg in soils, sunflower capitulums and sunflower pollen, respectively (Bonmatin et al., 2005). However, maximum imidacloprid residue concentrations collated from various applications scenarios and across multiple flowering plants can go up to concentrations of 6.59 × 10^3^ µg/kg in nectar (Zioga et al., 2020). This is just a little less than half of the concentration (1.59 × 10^4^ µg/L) at which the survival probability of *Eristalis* begins to significantly differ from the control group and therefore could induce both lethal and sublethal impacts in the fly. Hence, *Eristalis* may still be significantly affected in the field depending on the application method and its associated residue concentrations.

### 4.2 Implications of altered feeding times due to imidacloprid

Our observation that feeding times shorten across feeding sessions suggests that untreated flies learn and remember the nature of the feeder, providing some insight into cognitive and sensorimotor contributions to feeding efficiency. Feeding behaviour in our experiments can be split into localising and navigating to the food source and extracting the food from the capillary. In bees, locating food sources is typically improved by associating odours, colours, and landmark features to rewarding sites in a cognitive process which relies on an imperfect daily memory (Keasar et al., 1996; Raine et al., 2006; Evans et al., 2021). The motor skills required to extract food from a rewarding site can also be learnt and improved over time (Raine and Chittka, 2007). It is known that *Eristalis* makes use of floral visual cues to guide it to rewarding sites with very fixed innate colour preferences for yellow which triggers the proboscis extension reflex in this species (Dinkel and Lunau, 2001; Lunau et al., 2018).

Previous studies demonstrate that sublethal doses of imidacloprid can impair both cognitive and motor skills required for efficient foraging through the disruption of neural signals (Belzunces et al., 2012; Palmer et al., 2013; Mengoni Goñalons and Farina, 2018; Phelps et al., 2018; Li et al., 2019). Our data support a clear reduction in feeding efficiency at imidacloprid concentrations of 1.59×10^4^ µg/L, in terms of both longer time to reach the target ingested volume and a larger fraction of flies that fail to reach this target at all (Figure 4,5). We observed no such effects on feeding efficiency at lower doses, with both the feeding time (Figure 4) and the fraction of flies observed to utilise the feeder (figure 2) being similar to controls. While it remains difficult to determine whether reduced efficiency at high concentrations is due to both impaired cognitive functions and motor skills it is important to note that the1.59×10^4^ µg/L treatment was above our estimated chronic LD50 value, and well above the severe motor impairment ED50 value (Figure 7). Hence it seems likely that severe motor impairment in the fraction of surviving flies still able to feed at all after the first treatment at higher doses contributes strongly to their reduced efficiency.

### 4.3 Imidacloprid tolerance in *Eristalis*

Since survival rates remained above 50% for most concentrations after 72hrs (Fig. 7 B), we were unable to compute a reliable LD50 for acute exposure to imidacloprid. However, mortality rates did approach 50% at the highest median dose of 63.93 ng/mg. In comparison, estimates for honeybee LD50s 48hrs after ingestion are two to three orders of magnitudes lower, ranging from 0.04ng/mg to 0.67ng/mg (Suchail et al., 2000; Nauen et al., 2001). Our data are more consistent with another dipteran, *Drosophila melanogaster* which has an LD50 of 186-305ng/mg 8 days after acute exposure (Charpentier et al., 2014). The high tolerance of *D. melanogaster* and the sensitivity of honeybees are both attributed in parts to the interactions between detoxifying enzymes (p450 cytochromes) and imidacloprid. *D. melanogaster* overexpresses the CYP6G1 gene from the CYP6 family of p450 cytochromes which has also been repeatedly associated with high imidacloprid tolerance in the housefly, the brown planthopper, the sweet potato whitefly (Markussen and Kristensen, 2010; Ding et al., 2013; Yang et al., 2013). In contrast, the honeybee and the bumblebee (both relatively more sensitive to imidacloprid) express the CYP9Q3 and CYP9Q4 genes which result in a weak interaction between their detoxifying enzymes and nitro-substituted neonicotinoids such as imidacloprid (Manjon et al., 2018). The relatively high tolerance of *Eristalis* compared to honeybees could thus be attributed to the expression of CYP6 family genes and be part of a broader tolerance for pesticide in dipterans and other pests relative to honeybees. In addition, variability in pesticide sensitivity across species can also be affected by differences in the binding affinity of the pesticide to its target receptor and the subsequent signalling cascades (Bantz et al., 2018).

Aside from the family of CYP genes that *Eristalis* expresses, the interaction of these CYP genes with phytochemicals found in honey and pollen could be an alternative explanation for the elevated tolerance to imidacloprid observed in this species (Berenbaum and Johnson, 2015). *p*-Coumaric acid is one such phytochemical found in pollen and honey, both of which were used as dietary supplements throughout our experiments. *p*-Coumaric acid is known to broadly up-regulate detoxifying enzymes and in the case of the honeybee enhances the metabolism of Coumaphos, an organophosphate pesticide, in the midgut by approximately 60% (Mao et al., 2013). However, this phytochemical enhanced metabolism of imidacloprid does not appear as effective in honeybees, perhaps due to the naturally weak interactions between CYP9Q3 enzymes and imidacloprid (Manjon et al., 2018).

Although we observed no significant differences in survival or impairment between exposure to imidacloprid concentrations below 1.59 × 10^4^ µg/L and control, delayed effects occurring after the 18-day monitoring period cannot be excluded. Even acute exposure to pesticides have been shown to produce delayed effects on mortality rates, feeding, reproduction and foraging behaviour (Beketov and Liess, 2008; Wolz et al., 2021; Barascou et al., 2022). Although effects on mortality tend to happen within a week (Beketov and Liess, 2008), other more subtle effects like reduced foraging behaviour can only be observed almost two weeks after exposure (Barascou et al., 2022). While our long monitoring period provides some insurance against missing obvious delayed effects on mortality, our measures of locomotive impairment and feeding behaviours will not be able to detect more subtle sublethal effects including processes which occur at the cellular level in very tolerant species such as *D. melanogaster* (Martelli et al., 2020). This highlights the need to pair toxicological studies with more refined measures of behaviour which allow the early characterisation of various subtle sublethal effects.

### 4.4 Limitations of the current study and future research direction

Differences in male and female physiology and territoriality (Nordström et al., 2008) are likely to produce differences in imidacloprid impacts between sexes. Previous studies in dipterans suggest that *Eristalis* females may be more tolerant of imidacloprid than males (Charpentier et al., 2014; Kavi et al., 2014). While our choice to focus only on males in this study limits our ability to predict impacts on females it also allowed us to characterise the effect of imidacloprid more thoroughly on males, for which, the sensory ecology and visually mediated behaviours are better understood. Male *Eristalis* are aggressively territorial and will defend patches of flowers by hovering over an area and chasing other hoverflies and insects, both of which are visually demanding (Wellington and Fitzpatrick, 1981). The high visual sensitivity and large eye size in males has led to the adoption of male *Eristalis* as a major model for the study of visual processing of moving objects and patterns (Straw et al., 2006; Wiederman et al., 2007). Our thorough study of imidacloprid toxicity in male *Eristalis* here provides the foundation for future research on the impacts of imidacloprid on visually mediated foraging behaviour (Rigosi and O’Carroll, 2021).

Neither significant mortality nor locomotive impairment were observed until imidacloprid dosages exceeded the LD50 or ED50 respectively. Thereafter, the onset of impairment was fast and closely linked to high mortality. The relatively rapid onset of lethality at high imidacloprid dosages suggest that the current bioassay may be missing more subtle sublethal impacts such as the motor and cognitive deficits observed in other species (Decourtye et al., 2004; Parkinson and Gray, 2019). Although our experiments provide a precise quantification of imidacloprid intake and its temporal dynamics, detecting sublethal impacts on cognition and foraging behaviour seems to require a more sophisticated approach where the animal is allowed to move freely. Advanced tracking with real-time stimuli for freely moving animals is already available as open-source software (Stowers et al., 2017) and can be easily adapted to investigate the sublethal impacts of imidacloprid on foraging behaviour in *Eristalis*. Our study provides precise LD50 and ED50 for acute and chronic exposure scenarios which can be used to carefully select the appropriate dosages for these refined experiments. The combination of traditional toxicological assays with novel neuroethological techniques promises to be a powerful tool for research into the sublethal impacts of pesticides in the future.

## 5 Acknowledgements

We would like to acknowledge Johan Ahlgren for experimental assistance, and POLYFLY SL (Almeria, Spain) for the fast supply of drone fly pupae throughout the project.

## References

Bakker, L., van der Werf, W., Tittonell, P., Wyckhuys, K.A., and Bianchi, F.J. (2020). Neonicotinoids in global agriculture: evidence for a new pesticide treadmill? Ecology and Society 25(3).

Bantz, A., Camon, J., Froger, J.-A., Goven, D., and Raymond, V. (2018). Exposure to sublethal doses of insecticide and their effects on insects at cellular and physiological levels. Current opinion in insect science 30, 73–78.

Barascou, L., Requier, F., Sené, D., Crauser, D., Le Conte, Y., and Alaux, C. (2022). Delayed effects of a single dose of a neurotoxic pesticide (sulfoxaflor) on honeybee foraging activity. Science of The Total Environment 805, 150351.

Basley, K., Davenport, B., Vogiatzis, K., and Goulson, D. (2018). Effects of chronic exposure to thiamethoxam on larvae of the hoverfly Eristalis tenax (Diptera, Syrphidae). PeerJ 6, e4258.

Baumann, K., Keune, J., Wolters, V., and Jauker, F. (2021). Distribution and pollination services of wild bees and hoverflies along an altitudinal gradient in mountain hay meadows. Ecology and evolution 11(16), 11345–11351.

Beketov, M.A., and Liess, M. (2008). Acute and delayed effects of the neonicotinoid insecticide thiacloprid on seven freshwater arthropods. Environmental Toxicology and Chemistry 27(2), 461–470. doi: https://doi.org/10.1897/07-322R.1.

Belzunces, L.P., Tchamitchian, S., and Brunet, J.-L. (2012). Neural effects of insecticides in the honey bee. Apidologie 43(3), 348–370.

Berenbaum, M.R., and Johnson, R.M. (2015). Xenobiotic detoxification pathways in honey bees. Current opinion in insect science 10, 51–58.

Blümel, S., Bakker, F., Baier, B., Brown, K., Candolfi, M., Goßmann, A., et al. (2000). Laboratory residual contact test with the predatory mite Typhlodromus pyri Scheuten (Acari: Phytoseiidae) for regulatory testing of plant protection products. Guidelines to evaluate side-effects of plant protection products to non-target arthropods, 121–133.

Bonmatin, J.M., Moineau, I., Charvet, R., Colin, M.E., Fleche, C., and Bengsch, E.R. (2005). “Behaviour of Imidacloprid in Fields. Toxicity for Honey Bees,” in Environmental Chemistry: Green Chemistry and Pollutants in Ecosystems, eds. E. Lichtfouse, J. Schwarzbauer & D. Robert. (Berlin, Heidelberg: Springer Berlin Heidelberg), 483–494.

Cardillo, G. (2021). “Five parameters logistic regression - There and back again”. (Mathworks).

Cavallaro, M.C., Morrissey, C.A., Headley, J.V., Peru, K.M., and Liber, K. (2017). Comparative chronic toxicity of imidacloprid, clothianidin, and thiamethoxam to Chironomus dilutus and estimation of toxic equivalency factors. Environmental toxicology and chemistry 36(2), 372–382.

Charpentier, G.l., Louat, F., Bonmatin, J.-M., Marchand, P.A., Vanier, F., Locker, D., et al. (2014). Lethal and sublethal effects of imidacloprid, after chronic exposure, on the insect model Drosophila melanogaster. Environmental science & technology 48(7), 4096–4102.

Condalfi, M., SETAC-Europe, and EC (Year). “Guidance document on regulatory testing and risk assessment procedures for plant protection products with non-target arthropods”, in: ESCORT2: Society of Environment Toxicology and Chemistry).

Cook, D.F., Voss, S.C., Finch, J.T., Rader, R.C., Cook, J.M., and Spurr, C.J. (2020). The Role of Flies as Pollinators of Horticultural Crops: An Australian Case Study with Worldwide Relevance. Insects 11(6), 341. doi: 10.3390/insects11060341.

Cvetković, V.J., Mitrović, T.L., Jovanović, B., Stamenković, S.S., Todorović, M., Ðorđević, M., et al. (2015). Toxicity of dimethyl sulfoxide against Drosophila melanogaster. Biol Nyssana 6(2), 91–95.

Da Costa, L.M., Grella, T.C., Barbosa, R.A., Malaspina, O., and Nocelli, R.C.F. (2015). Determination of acute lethal doses (LD50 and LC50) of imidacloprid for the native bee Melipona scutellaris Latreille, 1811 (Hymenoptera: Apidae). Sociobiology 62(4), 578–582.

Decourtye, A., Devillers, J., Cluzeau, S., Charreton, M., and Pham-Delègue, M.-H. (2004). Effects of imidacloprid and deltamethrin on associative learning in honeybees under semi-field and laboratory conditions. Ecotoxicology and Environmental Safety 57(3), 410–419. doi: https://doi.org/10.1016/j.ecoenv.2003.08.001.

Decourtye, A., Lacassie, E., and Pham-Delègue, M.-H. (2003). Learning performances of honeybees (Apis mellifera L) are differentially affected by imidacloprid according to the season. Pest Management Science 59(3), 269–278. doi: https://doi.org/10.1002/ps.631.

Ding, Z., Wen, Y., Yang, B., Zhang, Y., Liu, S., Liu, Z., et al. (2013). Biochemical mechanisms of imidacloprid resistance in Nilaparvata lugens: Over-expression of cytochrome P450 CYP6AY1. Insect Biochemistry and Molecular Biology 43(11), 1021–1027. doi: https://doi.org/10.1016/j.ibmb.2013.08.005.

Dinkel, T., and Lunau, K. (2001). How drone flies (Eristalis tenax L., Syrphidae, Diptera) use floral guides to locate food sources. Journal of Insect Physiology 47(10), 1111–1118. doi: https://doi.org/10.1016/S0022-1910(01)00080-4.

Doyle, T., Hawkes, W.L., Massy, R., Powney, G.D., Menz, M.H., and Wotton, K.R. (2020). Pollination by hoverflies in the Anthropocene. Proceedings of the Royal Society B 287(1927), 20200508.

Durlak, J.A. (2009). How to select, calculate, and interpret effect sizes. Journal of pediatric psychology 34(9), 917–928.

Easton, A.H., and Goulson, D. (2013). The neonicotinoid insecticide imidacloprid repels pollinating flies and beetles at field-realistic concentrations. PLoS One 8(1), e54819.

Efron, B. (1982). The Jackknife, the Bootstrap, and Other Resampling Plans. Society for Industrial and Applied Mathematics.

EFSA (2013). Guidance on the risk assessment of plant protection products on bees (Apis mellifera, Bombus spp. and solitary bees). EFSA Journal 11(7), 3295.

EFSA (2015). Scientific Opinion addressing the state of the science on risk assessment of plant protection products for non-target arthropods. EFSA Journal 13(2), 3996.

EFSA (2018). Peer review of the pesticide risk assessment for bees for the active substance imidacloprid considering the uses as seed treatments and granules. EFSA Journal 16(2), e05178.

Evans, L.J., Smith, K.E., and Raine, N.E. (2021). Odour Learning Bees Have Longer Foraging Careers Than Non-learners in a Natural Environment. Frontiers in Ecology and Evolution 9(471). doi: 10.3389/fevo.2021.676289.

Gehan, E.A. (1965). A generalized Wilcoxon test for comparing arbitrarily singly-censored samples. Biometrika 52(1-2), 203–224.

Geiger, F., Bengtsson, J., Berendse, F., Weisser, W.W., Emmerson, M., Morales, M.B., et al. (2010). Persistent negative effects of pesticides on biodiversity and biological control potential on European farmland. Basic and Applied Ecology 11(2), 97–105.

Gervais, A., Chagnon, M., and Fournier, V. (2018). Diversity and pollen loads of flower flies (Diptera: Syrphidae) in cranberry crops. Annals of the Entomological Society of America 111(6), 326–334.

Gilbert, F. (1985). Diurnal activity patterns in hoverflies (Diptera: Syrphidae). Ecological entomology.

Grimm, C., Schmidli, H., Bakker, F., Brown, K., Campbell, P., Candolfi, M., et al. (2001). Use of standard toxicity tests with Typhlodromus pyri and Aphidius rhopalosiphi to establish a dose-response relationship. Anzeiger für Schädlingskunde= Journal of pest science 74(3), 72–84.

Hallmann, C.A., Sorg, M., Jongejans, E., Siepel, H., Hofland, N., Schwan, H., et al. (2017). More than 75 percent decline over 27 years in total flying insect biomass in protected areas. PLOS ONE 12(10), e0185809. doi: 10.1371/journal.pone.0185809.

He, Y., Zhao, J., Zheng, Y., Desneux, N., and Wu, K. (2012). Lethal effect of imidacloprid on the coccinellid predator Serangium japonicum and sublethal effects on predator voracity and on functional response to the whitefly Bemisia tabaci. Ecotoxicology 21(5), 1291–1300.

Hedges, L.V., Hedges, P.L.V., Olkin, I., and Olkin, j.a. (1985). Statistical Methods for Meta-Analysis. Elsevier Science.

Howlett, B.G., and Gee, M. (2019). The potential management of the drone fly (Eristalis tenax) as a crop pollinator in New Zealand. New Zealand Plant Protection 72, 221–230.

Jarlan, A., De Oliveira, D., and Gingras, J. (1997). Pollination by Eristalis tenax (Diptera: Syrphidae) and seed set of greenhouse sweet pepper. Journal of Economic Entomology 90(6), 1646–1649.

Kagabu, S. (2011). Discovery of Imidacloprid and Further Developments from Strategic Molecular Designs. Journal of Agricultural and Food Chemistry 59(7), 2887–2896. doi: 10.1021/jf101824y.

Kavi, L.A., Kaufman, P.E., and Scott, J.G. (2014). Genetics and mechanisms of imidacloprid resistance in house flies. Pesticide biochemistry and physiology 109, 64–69.

Keasar, T., Motro, U.Z.I., Shur, Y., and Shmida, A.V.I. (1996). Overnight memory retention of foraging skills by bumblebees is imperfect. Animal Behaviour 52(1), 95–104. doi: https://doi.org/10.1006/anbe.1996.0155.

Lagadic, L., Bernard, L., and Leicht, W. (1993). Topical and oral activities of imidacloprid and cyfluthrin against susceptible laboratory strains of Heliothis virescens and Spodoptera littoralis. Pesticide science 38(4), 323–328.

Laycock, I., and Cresswell, J.E. (2013). Repression and recuperation of brood production in Bombus terrestris bumble bees exposed to a pulse of the neonicotinoid pesticide imidacloprid. PloS one 8(11), e79872.

Li, Z., Yu, T., Chen, Y., Heerman, M., He, J., Huang, J., et al. (2019). Brain transcriptome of honey bees (Apis mellifera) exhibiting impaired olfactory learning induced by a sublethal dose of imidacloprid. Pesticide biochemistry and physiology 156, 36–43.

Lunau, K., An, L., Donda, M., Hohmann, M., Sermon, L., and Stegmanns, V. (2018). Limitations of learning in the proboscis reflex of the flower visiting syrphid fly Eristalis tenax. PLOS ONE 13(3), e0194167. doi: 10.1371/journal.pone.0194167.

Manjon, C., Troczka, B.J., Zaworra, M., Beadle, K., Randall, E., Hertlein, G., et al. (2018). Unravelling the Molecular Determinants of Bee Sensitivity to Neonicotinoid Insecticides. Current Biology 28(7), 1137–1143.e1135. doi: https://doi.org/10.1016/j.cub.2018.02.045.

Mao, W., Schuler, M.A., and Berenbaum, M.R. (2013). Honey constituents up-regulate detoxification and immunity genes in the western honey bee Apis mellifera. Proceedings of the National Academy of Sciences 110(22), 8842–8846.

Markussen, M.D.K., and Kristensen, M. (2010). Cytochrome P450 monooxygenase-mediated neonicotinoid resistance in the house fly Musca domestica L. Pesticide Biochemistry and Physiology 98(1), 50–58. doi: https://doi.org/10.1016/j.pestbp.2010.04.012.

Marletto, F., Patetta, A., and Manino, A. (2003). Laboratory assessment of pesticide toxicity to bumblebees. Bulletin of insectology 56(1), 155–158.

Martelli, F., Zhongyuan, Z., Wang, J., Wong, C.-O., Karagas, N.E., Roessner, U., et al. (2020). Low doses of the neonicotinoid insecticide imidacloprid induce ROS triggering neurological and metabolic impairments in Drosophila. Proceedings of the National Academy of Sciences 117(41), 25840–25850.

Mead-Briggs, M., Brown, K., Candolfi, M., Coulson, M., Miles, M., Moll, M., et al. (2000). A laboratory test for evaluating the effects of plant protection products on the parasitic wasp, Aphidius rhopalosiphi (DeStephani-Perez)(Hymenoptera: Braconidae). Guidelines to evaluate side-effects of plant protection products to non-target arthropods, 13–25.

Mengoni Goñalons, C., and Farina, W.M. (2018). Impaired associative learning after chronic exposure to pesticides in young adult honey bees. Journal of Experimental Biology 221(7), jeb176644.

Mizell, R.F., and Sconyers, M.C. (1992). Toxicity of imidacloprid to selected arthropod predators in the laboratory. The Florida Entomologist 75(2), 277–280.

Nauen, R. (1995). Behaviour modifying effects of low systemic concentrations of imidacloprid on Myzus persicae with special reference to an antifeeding response. Pesticide Science 44(2), 145–153. doi: https://doi.org/10.1002/ps.2780440207.

Nauen, R., Ebbinghaus-Kintscher, U., and Schmuck, R. (2001). Toxicity and nicotinic acetylcholine receptor interaction of imidacloprid and its metabolites in Apis mellifera (Hymenoptera: Apidae). Pest Management Science 57(7), 577–586. doi: https://doi.org/10.1002/ps.331.

Nicholas, S., Thyselius, M., Holden, M., and Nordström, K. (2018). Rearing and long-term maintenance of Eristalis tenax hoverflies for research studies. Journal of visualized experiments: JoVE (135).

Nordström, K., Barnett, P.D., Moyer de Miguel, I.M., Brinkworth, R.S.A., and O’Carroll, D.C. (2008). Sexual Dimorphism in the Hoverfly Motion Vision Pathway. Current Biology 18(9), 661–667. doi: https://doi.org/10.1016/j.cub.2008.03.061.

Ødegaard, F. (2008). How many species of arthropods? Erwin’s estimate revised. Biological Journal of the Linnean Society 71(4), 583–597. doi: 10.1111/j.1095-8312.2000.tb01279.x.

Overmyer, J., Mason, B., and Armbrust, K. (2005). Acute toxicity of imidacloprid and fipronil to a nontarget aquatic insect, Simulium vittatum Zetterstedt cytospecies IS-7. Bulletin of environmental contamination and toxicology 74(5), 872–879.

Palmer, M.J., Moffat, C., Saranzewa, N., Harvey, J., Wright, G.A., and Connolly, C.N. (2013). Cholinergic pesticides cause mushroom body neuronal inactivation in honeybees. Nature communications 4(1), 1–8.

Parkinson, R.H., and Gray, J.R. (2019). Neural conduction, visual motion detection, and insect flight behaviour are disrupted by low doses of imidacloprid and its metabolites. Neurotoxicology 72, 107–113.

Peto, R., and Peto, J. (1972). Asymptotically efficient rank invariant test procedures. Journal of the Royal Statistical Society: Series A (General) 135(2), 185–198.

Phelps, J.D., Strang, C.G., Gbylik-Sikorska, M., Sniegocki, T., Posyniak, A., and Sherry, D.F. (2018). Imidacloprid slows the development of preference for rewarding food sources in bumblebees (Bombus impatiens). Ecotoxicology 27(2), 175–187.

R Core Team (2021). “R: A language and program for sattistical computing”. (Vienna, Austria: R Foundation for statistical computing).

Raine, N.E., and Chittka, L. (2007). Pollen foraging: learning a complex motor skill by bumblebees (Bombus terrestris). Naturwissenschaften 94(6), 459–464.

Raine, N.E., Ings, T.C., Dornhaus, A., Saleh, N., and Chittka, L. (2006). Adaptation, genetic drift, pleiotropy, and history in the evolution of bee foraging behavior. Advances in the Study of Behavior 36, 305–354.

Rigosi, E., and O’Carroll, D.C. (2021). Acute application of imidacloprid alters the sensitivity of direction selective motion detecting neurons in an insect pollinator. Frontiers in Physiology 12, 960.

Sakamoto, H., and Goka, K. (2021). Acute toxicity of typical ant control agents to the red imported fire ant, Solenopsis invicta (Hymenoptera: Formicidae). Applied Entomology and Zoology 56(2), 217–224.

Sánchez-Bayo, F., and Wyckhuys, K.A. (2019). Worldwide decline of the entomofauna: A review of its drivers. Biological conservation 232, 8–27.

Simon-Delso, N., Amaral-Rogers, V., Belzunces, L.P., Bonmatin, J.M., Chagnon, M., Downs, C., et al. (2015). Systemic insecticides (neonicotinoids and fipronil): trends, uses, mode of action and metabolites. Environmental Science and Pollution Research 22(1), 5–34. doi: 10.1007/s11356-014-3470-y.

Soares, H.M., Jacob, C.R.O., Carvalho, S.M., Nocelli, R.C.F., and Malaspina, O. (2015). Toxicity of imidacloprid to the stingless bee Scaptotrigona postica Latreille, 1807 (Hymenoptera: Apidae). Bulletin of environmental contamination and toxicology 94(6), 675–680.

Ssymank, A., Kearns, C., Pape, T., and Thompson, F.C. (2008). Pollinating flies (Diptera): a major contribution to plant diversity and agricultural production. Biodiversity 9(1-2), 86–89.

Stoughton, S.J., Liber, K., Culp, J., and Cessna, A. (2008). Acute and chronic toxicity of imidacloprid to the aquatic invertebrates Chironomus tentans and Hyalella azteca under constant-and pulse-exposure conditions. Archives of Environmental Contamination and Toxicology 54(4), 662–673.

Stowers, J.R., Hofbauer, M., Bastien, R., Griessner, J., Higgins, P., Farooqui, S., et al. (2017). Virtual reality for freely moving animals. Nature Methods 14(10), 995–1002. doi: 10.1038/nmeth.4399.

Straw, A.D., Warrant, E.J., and O’Carroll, D.C. (2006). Abright zone’in male hoverfly (Eristalis tenax) eyes and associated faster motion detection and increased contrast sensitivity. Journal of Experimental Biology 209(21), 4339–4354.

Stubbs, A.E., and Falk, S.J. (1983). British Hoverflies: An Illustrated Identification Guide. British Entomological & Natural History Society.

Suchail, S., Guez, D., and Belzunces, L.P. (2000). Characteristics of imidacloprid toxicity in two Apis mellifera subspecies. Environmental Toxicology and Chemistry: An International Journal 19(7), 1901–1905.

Suchail, S., Guez, D., and Belzunces, L.P. (2001). Discrepancy between acute and chronic toxicity induced by imidacloprid and its metabolites in Apis mellifera. Environmental Toxicology and Chemistry: An International Journal 20(11), 2482–2486.

Tappert, L., Pokorny, T., Hofferberth, J., and Ruther, J. (2017). Sublethal doses of imidacloprid disrupt sexual communication and host finding in a parasitoid wasp. Scientific reports 7(1), 1–9.

Tharp, C.I., Johnson, G.D., and Onsager, J.A. (2000). Laboratory and field evaluations of imidacloprid against Melanoplus sanguinipes (Orthoptera: Acrididae) on small grains. Journal of economic entomology 93(2), 293–299.

Thompson, H.M., Wilkins, S., Harkin, S., Milner, S., and Walters, K.F. (2015). Neonicotinoids and bumblebees (Bombus terrestris): effects on nectar consumption in individual workers. Pest management science 71(7), 946–950.

Tomlin, C.D. (2006). “The pesticide manual: A world compendium.” (Surrey, UK: British Crop Protection Council), 598–599.

Wang, K.-Y., Liu, T.-X., Yu, C.-H., Jiang, X.-Y., and Yi, M.-Q. (2002). Resistance of Aphis gossypii (Homoptera: Aphididae) to fenvalerate and imidacloprid and activities of detoxification enzymes on cotton and cucumber. Journal of Economic Entomology 95(2), 407–413.

Wellington, W., and Fitzpatrick, S.M. (1981). Territoriality in the drone fly, Eristalis tenax (Diptera: Syrphidae). The Canadian Entomologist 113(8), 695–704.

Wiederman, S., Shoemaker, P., and O’carroll, D. (Year). “Biologically inspired small target detection mechanisms”, in: 2007 3rd International Conference on Intelligent Sensors, Sensor Networks and Information: IEEE), 269–273.

Wolz, M., Schrader, A., and Müller, C. (2021). Direct and delayed effects of exposure to a sublethal concentration of the insecticide λ-cyhalothrin on food consumption and reproduction of a leaf beetle. Science of The Total Environment 760, 143381. doi: https://doi.org/10.1016/j.scitotenv.2020.143381.

Yang, X., Xie, W., Wang, S.-l., Wu, Q.-j., Pan, H.-p., Li, R.-m., et al. (2013). Two cytochrome P450 genes are involved in imidacloprid resistance in field populations of the whitefly, Bemisia tabaci, in China. Pesticide Biochemistry and Physiology 107(3), 343–350. doi: https://doi.org/10.1016/j.pestbp.2013.10.002.

Zioga, E., Kelly, R., White, B., and Stout, J.C. (2020). Plant protection product residues in plant pollen and nectar: A review of current knowledge. Environmental research 189, 109873.

